# Maternal Biological Age Assessed in Early Pregnancy is Associated with Gestational Age at Birth

**DOI:** 10.1101/2021.01.11.425979

**Authors:** Eva E. Lancaster, Dana M. Lapato, Colleen Jackson-Cook, Jerome F. Strauss, Roxann Roberson-Nay, Timothy P. York

## Abstract

Maternal age is an established predictor of preterm birth independent of other recognized risk factors. The use of chronological age makes the assumption that individuals age at a similar rate. Therefore, it does not capture interindividual differences that may exist due to genetic background and environmental exposures. As a result, there is a need to identify biomarkers that more closely index the rate of cellular aging. One potential candidate is biological age (BA) estimated by the DNA methylome. This study investigated whether maternal BA, estimated in either early and/or late pregnancy, predicts gestational age at birth. BA was estimated from a genome-wide DNA methylation platform using the Horvath algorithm. Linear regression methods assessed the relationship between BA and pregnancy outcomes, including gestational age at birth and perceived stress during pregnancy, in a primary and replication cohort. Prenatal BA estimates from early pregnancy explained variance in gestational age at birth above and beyond the influence of other recognized preterm birth risk factors. Sensitivity analyses indicated that this signal was driven primarily by self-identified African American participants. This predictive relationship was sensitive to small variations in the BA estimation algorithm. Benefits and limitations of using BA in translational research and clinical applications for preterm birth are considered.

## 1 Introduction

Preterm birth (PTB; birth before 37 completed weeks of gestation) remains the leading contributor to neonatal mortality and morbidity worldwide. ^1^ In addition to the emotional distress associated with PTB, the monetary costs associated with PTB complications in the United States exceeded $28 billion dollars in 2005 alone. ^2^ Prenatal interventions to reduce the prevalence of PTB have shown promise, but identifying women at high risk for preterm delivery can be challenging. Results from epidemiological and family studies confirm that genetic, environmental, and behavioral factors all jointly influence PTB risk liability. ^3,4,5,6^ However, translating these results into improved clinical prediction models has proven difficult. One potential avenue for relating these three contributory sources to PTB risk is through DNA methylation-based biological age estimates.

Biological age (BA) describes the rate of cellular aging and progression towards senescence. Conventionally, researchers and clinicians have used chronological age to proxy BA, but accumulating evidence suggests that deviations between BA and chronological age are informative about risk for future adverse health outcomes, such as early mortality and cancer. ^7,8,9,10,11^ The notion that advanced BA indexes biological changes with respect to aging and senescence is supported by association studies with health outcomes, ^7^ including (but not limited to) reports that individuals with Werner and Hutchinson-Gilford progeria syndromes, genetic disorders of premature aging, exhibit markedly advanced BAs. ^12,13^ BA is most commonly estimated by patterns of DNA methylation (DNAm), which have been shown to correlate with chronological age. ^14^ DNAm is an epigenetic modification to DNA associated with genomic stability, transcriptional activity, and chromatin conformation. DNAm patterns change over time as a function of normal physiology. ^15^ Several hundred genomic loci have been robustly associated with age-related DNAm remodeling, and the DNAm levels at these sites are used to estimate BA. ^12,16,17^ The rationale that DNAm may index cellular aging stems from the susceptibility of DNAm remodeling to genetic, environmental, and behavioral factors which change throughout the life course. Moreover, aberrant DNAm patterns have been associated with negative health outcomes, including congenital disorders, developmental delay, and elevated risk for cancer, which further underscores the notion that unexpected changes in DNAm (and BA) are salient to current and future health outcomes. ^15^

Three primary lines of evidence support investigating the potential of BA to improve clinical predictive models of PTB risk. One, BA calculated from DNAm may reflect influences of past behaviors (e.g., smoking) and environmental exposures (e.g., pollution, trauma). ^18,19,20^ This sensitivity to PTB risk factors alone suggests that BA may be more useful than chronological age, which is uniform regardless of life experiences. Two, incorporating genomic information in the form of polygenic risk scores (PRS) has improved clinical prediction algorithms for other multifactorial disorders, like breast cancer, prostate cancer, and type 1 diabetes. ^21,22^ Similar success may be possible for cumulative epigenomic summaries like BA. Three, significant racial health disparities in PTB rates have persisted in the US for decades between individuals who self-identified as non-Hispanic African American (AA) and non-Hispanic European American (EA). ^23^ A putative driver of this disparity is biological weathering, which is premature cellular deterioration due to chronic social, economic, and environmental stressors. ^24,25,26^ One of the strongest pieces of evidence supporting the theory of biological weathering is the observation that advanced maternal age-related perinatal complications begin on average at younger ages for AA women compared to EA women. ^27^ BA provides a plausible biological mechanism to account for the weathering hypothesis. Moreover, the observed variability in which risk for age-related complications begins further underscores the idea that BA may be more informative of individual risk than chronological age.

The purpose of this study was to explore the relationship between BA and gestational age at delivery (GAAD) in a racially diverse longitudinal cohort of pregnant women. To date, PTB research has primarily focused on investigating postnatal fetal measurements of cellular aging rather than maternal BA during pregnancy. ^28,29,30,31,32^ By measuring fetal BA, these studies could be assessing epigenetic changes that provide information about the developmental maturity of an infant at birth, rather than biological processes related to the onset of labor. As a result, little is known about the behavior of maternal BA during pregnancy and the relationship between maternal BA and pregnancy outcomes. This study sought to address these gaps by using repeated measures of DNAm from two longitudinal cohorts of pregnant women to characterize the stability of maternal DNAm-based BA across pregnancy, assess its relationship with GAAD, and determine whether these predictions vary by self-identified Census-based race category. The impact of chronological age, prenatal perceived stress level and tobacco smoking on the association between BA and GAAD was considered to evaluate the potential for BA to account for additional variation in GAAD above and beyond these established PTB risk factors. BA stability during pregnancy also was examined to identify informative intervals for evaluating PTB risk. Replication analyses were conducted in an independent cohort.

## 2 Material and Methods

### 2.1 Study cohort

#### Pregnancy, Race, Environment, Genes (PREG)

The Pregnancy, Race, Environment, Genes (PREG) Study is a prospective longitudinal cohort assessing the relationship between epigenetic factors, environmental exposures, and pregnancy outcomes. ^33^ Self-report questionnaires and maternal peripheral blood samples were collected up to four times throughout pregnancy. Inclusion criteria at enrollment were 1) singleton pregnancy conceived without assisted reproductive technology, 2) mother was 18-40 years old with no diagnosis of diabetes, 3) enrollment before 24 completed weeks of gestation, 4) mother and father had to self-identify as either both White or both Black without Latinx or Middle Eastern ancestry. The rationale for limiting the cohort by ancestry was to maximize the statistical power for genetic/epigenetic analyses and to investigate the role of environmental and epigenetic factors to perinatal health disparities. Exclusion criteria included diagnosis of maternal blood pressure disorders (e.g., preeclampsia), fetal congenital anomalies, placental or amniotic anomalies (e.g., placenta previa, polyhydramnios), fewer than three study time points completed, or use of a cerclage. Gestational age (GA) was confirmed by ultrasound. GA at each study visit and GAAD were recorded in days since conception.

### 2.2 Replication cohort

#### Global Alliance to Prevent Prematurity and Stillbirth (GAPPS)

Maternal blood specimens were obtained from the Global Alliance to Prevent Prematurity and Stillbirth (GAPPS) BioServices repository. GAPPS participant selection criteria matched all PREG study inclusion and most exclusion criteria to facilitate cross-study comparisons. African American samples were not available from GAPPS at the time of study initiation. Maternal peripheral blood samples were collected along with self-report questionnaires up to three times across pregnancy. Due to the smaller number of total possible study visits, GAPPS participants were included if they had least two time points of data. GAAD was reported in days since conception, but GA at each study visit was reported as trimester (i.e., 1, 2, or 3).

### 2.3 Biological age measurement

BA was estimated from genome-wide DNAm measurements using the Horvath method. ^12^ The Horvath algorithm calculates BA from DNAm levels at 353 genomic loci each measured by a single probe. Most of the loci only contribute modestly to the final age estimate (i.e., median weight is 6 weeks; range is 0.00000594 to 3.07 years). ^12^ Both PREG and GAPPS measured DNAm from peripheral blood specimens using Illumina microarray technology. The PREG study used the Infinium HumanMethylation450 BeadChip (450k); GAPPS, the Infinium EPIC BeadChip (850k). The 850k array is a newer sister technology to the 450k and includes 92% of the 450k probe set. The newer 850k array design omits 17 of the Horvath probes (4.8%). Despite the probe set differences, previous reports have suggested that the Horvath age estimates are only slightly underestimated in peripheral blood when these probes are missing (r *>* 0.91, n = 172). ^34,35^ Both PREG and GAPPS microarray experiments were separately performed at HudsonAlpha Institute for Biotechnology according to the manufacturer’s protocol (Illumina, San Diego, CA, USA). For both cohorts, the individual specimen placement were randomized on the array, but all specimens from a single participant were loaded onto a single array to minimize potential batch effects (see supplemental methods).

Before calculating BA, the quality of DNAm microarrays was assessed (Figure 1) using the Bioconductor R package minfi. ^36^ Probes with either poor signal intensity or known cross-hybridization activity were removed in accordance with established best practices (see supplement for additional details). Principal components analysis was used to identify potential experimental artifacts (e.g., batch effects), and based on this analysis, probe Beta-values were adjusted for positional effects using ComBat. ^37^ BA estimates for each specimen were calculated from adjusted Beta-values using the wateRmelon R package. ^38^ All statistical analyses were conducted in the R environment (version 3.5) ^39^.

**Figure 1.**
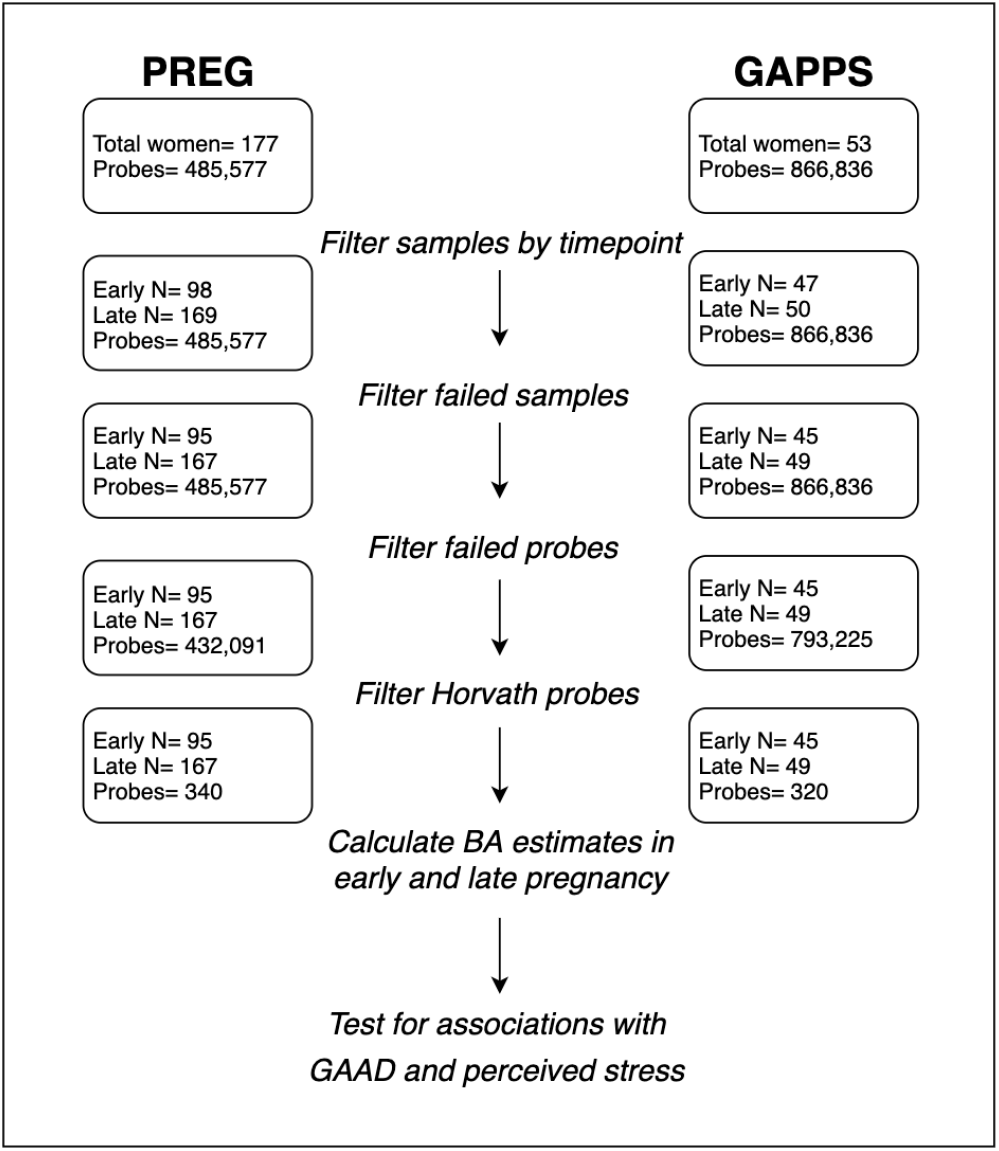
Diagram of study design and processing steps, highlighting probes and samples remaining in the primary (PREG) and replication (GAPPS) cohorts after performing quality control for DNA methylation microarrays. **Abbreviations**. BA = Horvath-derived biological age estimates; GAAD = gestational age at delivery.

### 2.4 Perceived stress measurement

The Perceived Stress Scale (PSS) is a ten question validated self-report instrument for assessing the magnitude and severity of recent stress levels. ^40^ Each item is a 5-point Likert-type question, with 0 indicating “never” and 4 indicating “very often”. Possible scores range from 0 to 40 with higher scores indicating greater levels and interference of perceived stress. The PSS was administered at every visit for the PREG study and in the second and third trimester health questionnaires for the GAPPS study.

### 2.5 Data analysis

Linear regression was used to test the relationship between BA estimates from early and late pregnancy with GAAD and prenatal perceived stress. To harmonize the data across studies while maintaining sample size, early and late prenatal DNAm measurements were defined in PREG as blood specimen obtained at a GA less than 100 days and after 180 days, respectively. In GAPPS, early pregnancy was defined as measurements collected in the first trimester while late pregnancy measurements were those obtained in the third trimester. To control for individual differences in chronological age, maternal age (collected at the time of study enrollment), was included as a covariate in all analyses. Lifetime smoking status (i.e., never, former, current), self-reported race, and prenatal perceived stress levels were included as covariates in the regression models for sensitivity analyses. Cell-type proportion estimates were not included because the Horvath BA algorithm is robust to biases related to cell-type heterogeneity. ^12^ Prenatal BA trajectories were characterized using linear latent growth curve models evaluated in Mplus and built using the R package MplusAutomation. ^41^ The purpose of the growth curve model was to quantify the interindividual difference in the baseline and rate of change of BA estimates across pregnancy.

## 3 Results

### 3.1 Participant demographics

After filtering for all inclusion/exclusion criteria, the PREG cohort consisted of 177 women who self-identified as Black (n = 89) or non-Hispanic White not of Middle Eastern descent (n = 88). Meeting the same criteria, the GAPPS cohort included 52 women who all self-identified as non-Hispanic Caucasian and not as African American (see Table 1 for additional participant demographics). In order to maintain consistency across cohorts, European American (EA) and African American (AA) will be used to describe women who self-identified as non-Hispanic White/Caucasian and Black/African American, respectively. The demographic attributes of the PREG EA subset and GAPPS cohort were on average more similar to each other than to the PREG AA subset. Overall, PREG AA women were more likely to be younger, report higher levels of perceived stress, and were less likely to report taking daily prenatal vitamins. The PTB rate for PREG and GAPPS cohorts were similar (PREG = 5.1%, GAPPS = 5.8%), but PREG AA women had significantly earlier GAAD. BA estimates were nominally higher than chronological age (Table 1 and Figure 4). Maternal chronological age and BA was moderately correlated in the PREG study (Pearson’s; 0.63 and 0.74 [EA = 0.42 and 0.62, AA = 0.67 and 0.73], in early and late pregnancy, respectively). The correlation between chronological age and BA was 0.71 in early pregnancy and 0.66 in late pregnancy (Pearson’s) in the GAPPS cohort.

**Table 1:**
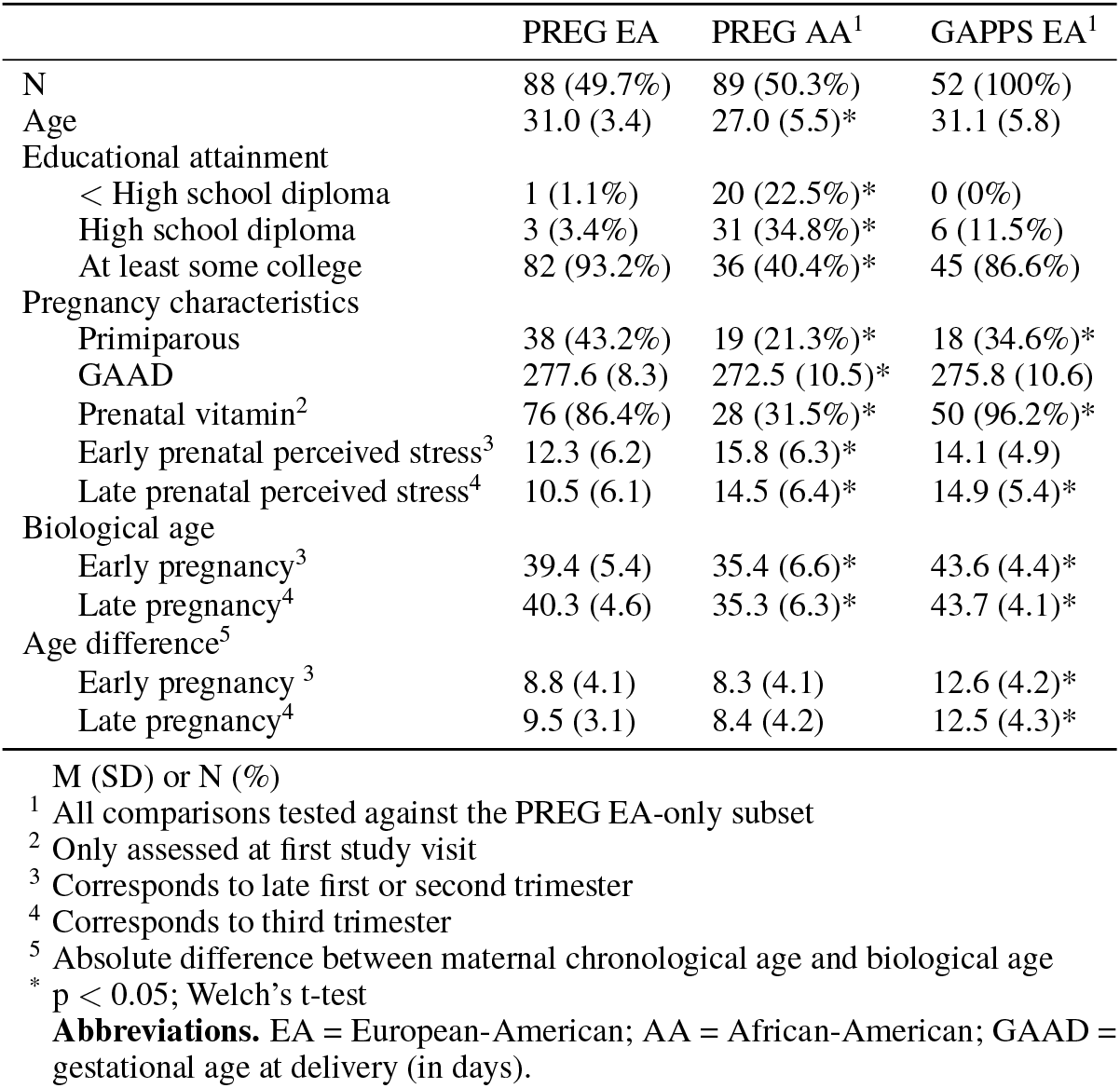
Cohort characteristics

|  | PREG EA | PREG AA<sup>1</sup> | GAPPS EA<sup>1</sup> |
| --- | --- | --- | --- |
| N | 88 (49.7%) | 89 (50.3%) | 52 (100%) |
| Age | 31.0 (3.4) | 27.0 (5.5)* | 31.1 (5.8) |
| Educational attainment |  |  |  |
| < High school diploma | 1 (1.1%) | 20 (22.5%)* | 0 (0%) |
| High school diploma | 3 (3.4%) | 31 (34.8%)* | 6 (11.5%) |
| At least some college | 82 (93.2%) | 36 (40.4%)* | 45 (86.6%) |
| Pregnancy characteristics |  |  |  |
| Primiparous | 38 (43.2%) | 19 (21.3%)* | 18 (34.6%)* |
| GAAD | 277.6 (8.3) | 272.5 (10.5)* | 275.8 (10.6) |
| Prenatal vitamin<sup>2</sup> | 76 (86.4%) | 28 (31.5%)* | 50 (96.2%)* |
| Early prenatal perceived stress<sup>3</sup> | 12.3 (6.2) | 15.8 (6.3)* | 14.1 (4.9) |
| Late prenatal perceived stress<sup>4</sup> | 10.5 (6.1) | 14.5 (6.4)* | 14.9 (5.4)* |
| Biological age |  |  |  |
| Early pregnancy<sup>3</sup> | 39.4 (5.4) | 35.4 (6.6)* | 43.6 (4.4)* |
| Late pregnancy<sup>4</sup> | 40.3 (4.6) | 35.3 (6.3)* | 43.7 (4.1)* |
| Age difference<sup>5</sup> |  |  |  |
| Early pregnancy<sup>3</sup> | 8.8 (4.1) | 8.3 (4.1) | 12.6 (4.2)* |
| Late pregnancy<sup>4</sup> | 9.5 (3.1) | 8.4 (4.2) | 12.5 (4.3)* |

M (SD) or N (%)
<sup>1</sup> All comparisons tested against the PREG EA-only subset
<sup>2</sup> Only assessed at first study visit
<sup>3</sup> Corresponds to late first or second trimester
<sup>4</sup> Corresponds to third trimester
<sup>5</sup> Absolute difference between maternal chronological age and biological age
\* $p < 0.05$ ; Welch's t-test
**Abbreviations.** EA = European-American; AA = African-American; GAAD = gestational age at delivery (in days).

**Figure 4.**
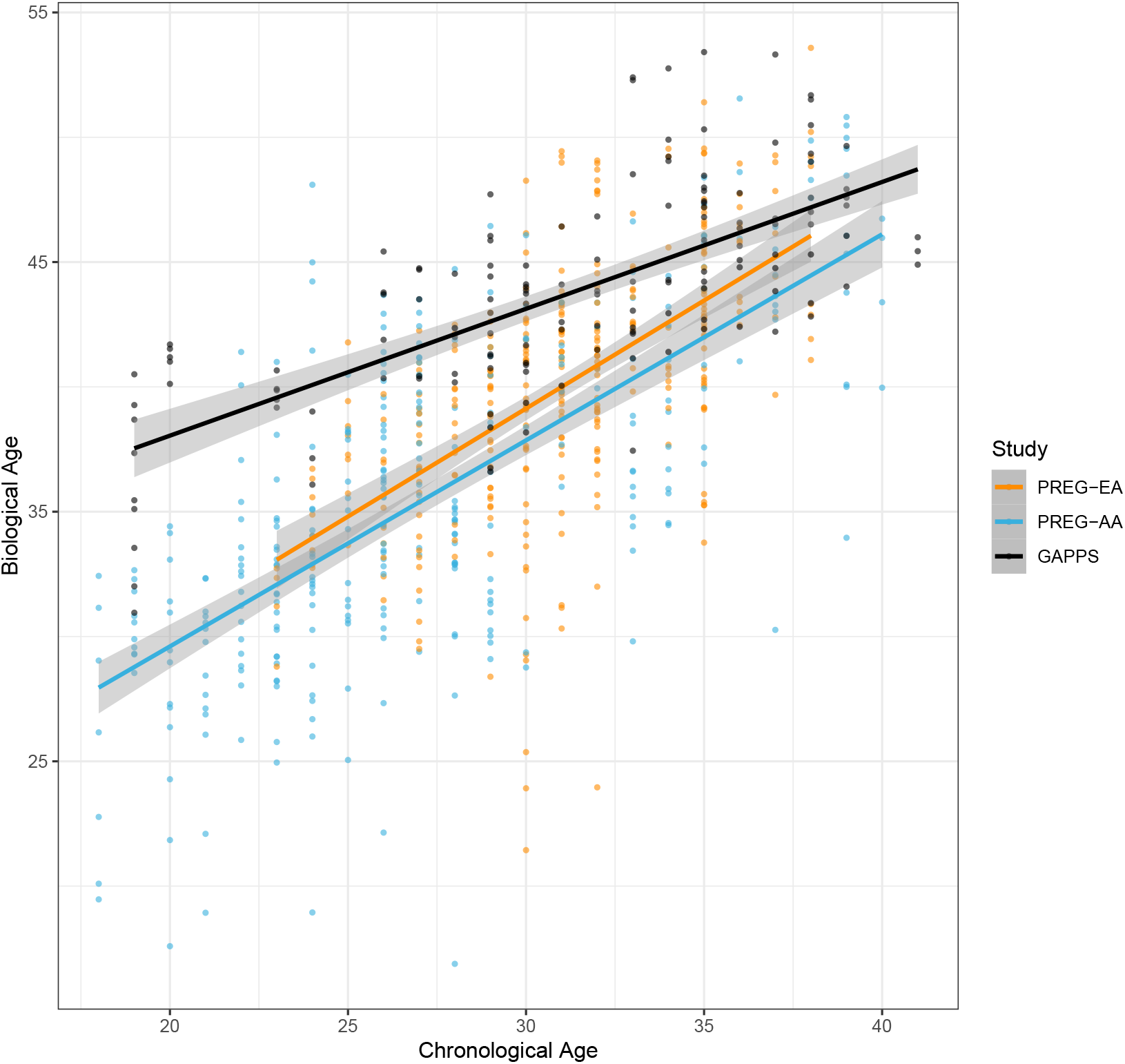
Correlation between chronological age and DNA methylation-based biological age estimates. The variance explained by chronological age and Horvath biological age estimates is similar between the self-identified European American (EA) and African American (AA) groups in the PREG study but different across studies.

After filtering DNAm data based on quality metrics, 262 and 94 person time points of data remained for the PREG and GAPPS cohorts, respectively. Subsequent division of measures based on GA at assessment resulted in 95 early pregnancy time points (EA = 49, AA = 46) and 167 late pregnancy time points (EA = 85, AA = 82) in PREG (Figure 1). The GAPPS cohort consisted of 45 early pregnancy measurements and 49 late pregnancy measurements. During preprocessing steps, 13 of the Horvath probes were identified as poor quality and removed from PREG, and 33 probes removed in GAPPS (46 total unique Horvath probes between both cohorts). To assess the impact of different probe subsets, analyses were performed with both the largest possible Horvath probe set for each cohort (PREG = 340 [96%], GAPPS = 320 [91%] (see Figure 1) and with the subset of Horvath probes shared in common between the two cohorts (n = 307 [87%]).

### 3.2 Association between BA and GAAD

The coefficients, standard errors, and p-values for all models tested with the PREG cohort are reported in Table 2. For each model, BA and GAAD were the predictor and response variables, respectively. In the full PREG sample, BA estimates outperformed chronological age in predicting GAAD (adjusted R-squared = 7.67% and 3.57%, respectively). The full PREG sample showed a significant relationship between the early pregnancy Horvath-derived BA estimates and GAAD (p-value threshold < 0.008 after Bonferroni adjustment for multiple testing). Higher BA estimates had a positive relationship with GAAD, indicating that an earlier GAAD is associated with younger BAs. Although the relationship between BA and GAAD was primarily supported by the AA subset, the significant relationship between BA and GAAD in the full sample remained after including a self-reported race variable in the model (p = 0.006). However, the relationship between early prenatal BA and GAAD was attenuated when retaining the maximum number of probes available (p = 0.005 in n = 340 probes [Supplementary Table S1]; p = 0.003 in n = 320 probes [Table 2]). There were no significant findings between GAAD and late pregnancy BA estimates.

**Table 2:** Relationships between Gestational Age at Delivery, Perceived Stress, and Biological Age Estimates in the PREG cohort^12^

|  | Full Sample<br>Coef | Full Sample<br>SE | Full Sample<br>P-value | EA Subset<br>Coef | EA Subset<br>SE | EA Subset<br>P-value | AA Subset<br>Coef | AA Subset<br>SE | AA Subset<br>P-value |
| --- | --- | --- | --- | --- | --- | --- | --- | --- | --- |
| Predicts GAAD |  |  |  |  |  |  |  |  |  |
| early BA | 0.63 | 0.21 | 0.003* | 0.40 | 0.24 | 0.107 | 0.71 | 0.33 | 0.038 |
| late BA | 0.06 | 0.18 | 0.763 | -0.11 | 0.26 | 0.669 | 0.03 | 0.27 | 0.918 |
| Predicts early prenatal PSS |  |  |  |  |  |  |  |  |  |
| early BA | -0.35 | 0.13 | 0.009 | -0.16 | 0.19 | 0.389 | -0.43 | 0.17 | 0.016 |
| Predicts late prenatal PSS |  |  |  |  |  |  |  |  |  |
| early BA | -0.14 | 0.14 | 0.319 | -0.11 | 0.21 | 0.600 | -0.15 | 0.19 | 0.435 |
| late BA | -0.09 | 0.13 | 0.514 | -0.07 | 0.20 | 0.715 | 0.07 | 0.17 | 0.669 |
| Predicts late BA |  |  |  |  |  |  |  |  |  |
| early PSS | -0.08 | 0.05 | 0.098 | -0.06 | 0.06 | 0.365 | -0.03 | 0.08 | 0.742 |

<sup>1</sup> Horvath probe sets were reduced to match the probes available for GAPPS.
<sup>2</sup> Maternal chronological age was included as a covariate in all models.
\* Survives Bonferroni adjustment for 6 tests, $P\text{-val} < 0.008$
**Abbreviations.** Coef = coefficient; SE = standard error; EA = European-American; AA = African-American; PSS = perceived stress scale total score; GAAD = gestational age at delivery; BA = Horvath-derived biological age estimates.

**Table 3:** Relationships between Gestational Age at Delivery, Perceived Stress, and Biological Age Estimates in the GAPPS replication cohort ^1,2^

|  | Coef | SE | P-value |
| --- | --- | --- | --- |
| Predicts GAAD |  |  |  |
| early BA | -0.11 | 0.50 | 0.821 |
| late BA | 0.37 | 0.41 | 0.371 |
| Predicts early prenatal PSS |  |  |  |
| early BA | -0.23 | 0.23 | 0.341 |
| Predicts late prenatal PSS |  |  |  |
| early BA | -0.40 | 0.27 | 0.139 |
| late BA | -0.14 | 0.26 | 0.599 |
| Predicts late BA |  |  |  |
| early PSS | -0.01 | 0.10 | 0.898 |

<sup>1</sup> Horvath probe sets were reduced to match the probes available for PREG.
<sup>2</sup> Maternal chronological age was included as a covariate in all models.
\* Survives Bonferroni adjustment $P\text{-val} < 0.008$
**Abbreviations.** Coef = coefficient; SE = standard error; PSS = perceived stress scale total score; GAAD = gestational age at delivery; BA = Horvath-derived biological age estimates.

A marginally significant relationship between prenatal PSS and BA estimates in early pregnancy was identified (p = 0.009) in the full sample. Similar to the direction of the relationship identified in the GAAD analyses, a higher PSS was associated with a lower BA. A nominally significant relationship between BA and GAAD remained even after adjusting for perceived stress in early pregnancy (p = 0.012). A follow-up analysis in the GAPPS sample, composed entirely of women with EA ancestry, showed no significant relationships between BA and GAAD or between perceived stress and BA.

### 3.3 Evaluation of BA as a potential clinical marker for GAAD

Residualized BA scores were calculated by regressing BA onto chronological age and reflect the deviation between chronological age and BA. For PREG, residualized BA scores were calculated using the largest possible Horvath probe subset (n = 340). Overall, BA residualized scores were relatively stable over the course of pregnancy regardless of self-identified race and had significant between-person heterogeneity (Figure 2). A significant relationship between BA baseline measurement (i.e., the model intercept), but not rate of change across pregnancy (i.e., the slope of the model), and GAAD was identified (see supplement). This finding is largely in agreement with the results from the linear regression models showing early BA associated with GAAD.

**Figure 2.**
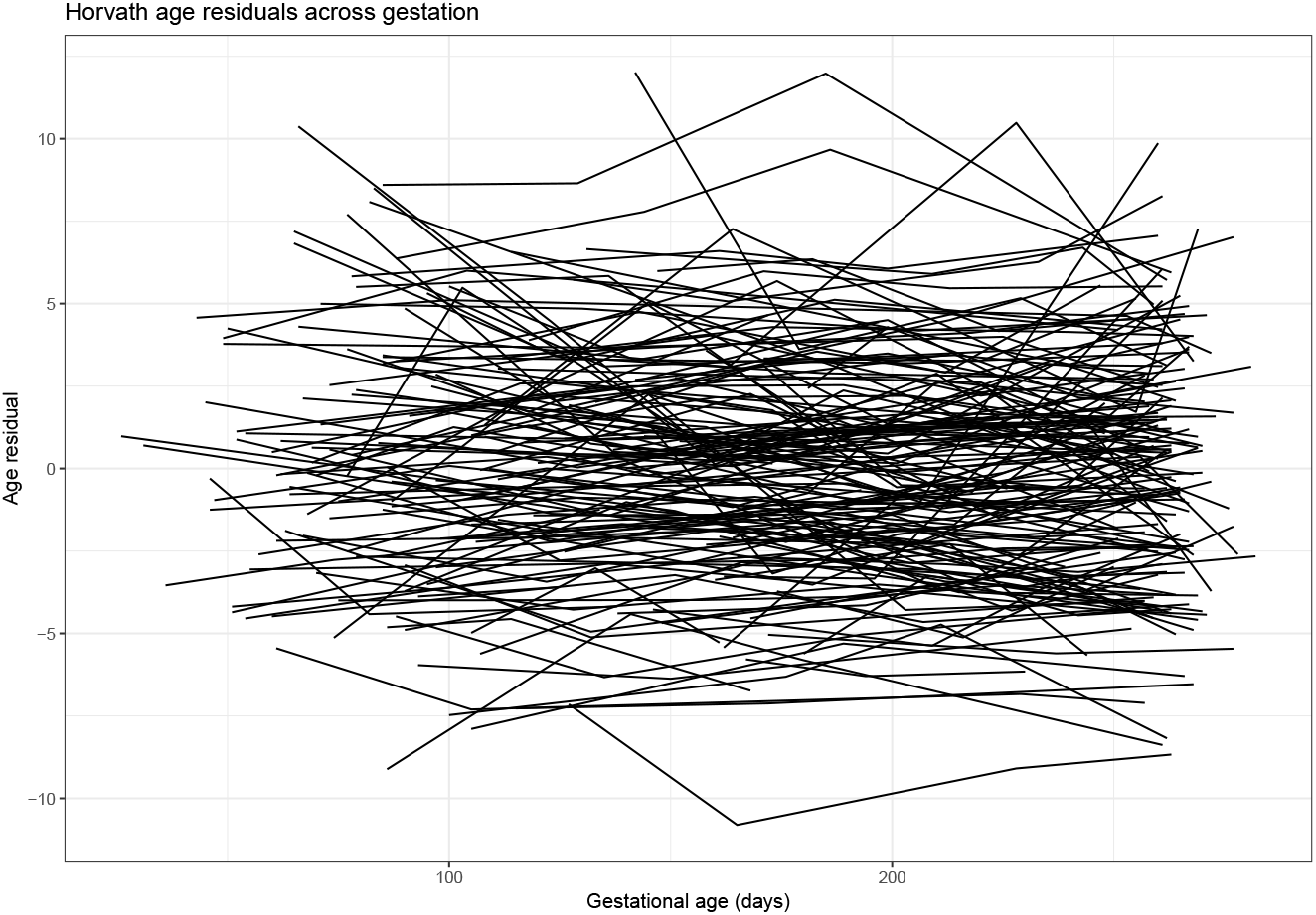
Biological age residuals across pregnancy. Residualized biological age scores were calculated for the entire PREG sample by regressing biological age estimates on to chronological age. Each line represents an unique individual, and each point indicates an individual assessment (i.e., study visit).

Critically, there was greater variability in the residualized BA scores in the PREG AA subset compared to the EA subset (Figure 3a). Follow up analyses revealed that BA residualized scores were sensitive to probe subset size and self-identified race. Residualized scores were calculated for both the full PREG (n = 340) and shared (n = 307) Horvath probe subsets, and self-identified Census-based race significantly predicted BA residuals for the shared probe set above and beyond the BA residuals for the full PREG BA subset (t-value = −5.89; Figure 3b). The sensitivity of BA estimation to probe subset size and composition was further highlighted by comparing the correlation between BA and chronological age in the PREG and GAPPS cohorts, which had different subsets of Horvath probes available (Figure 4).

**Figure 3.**
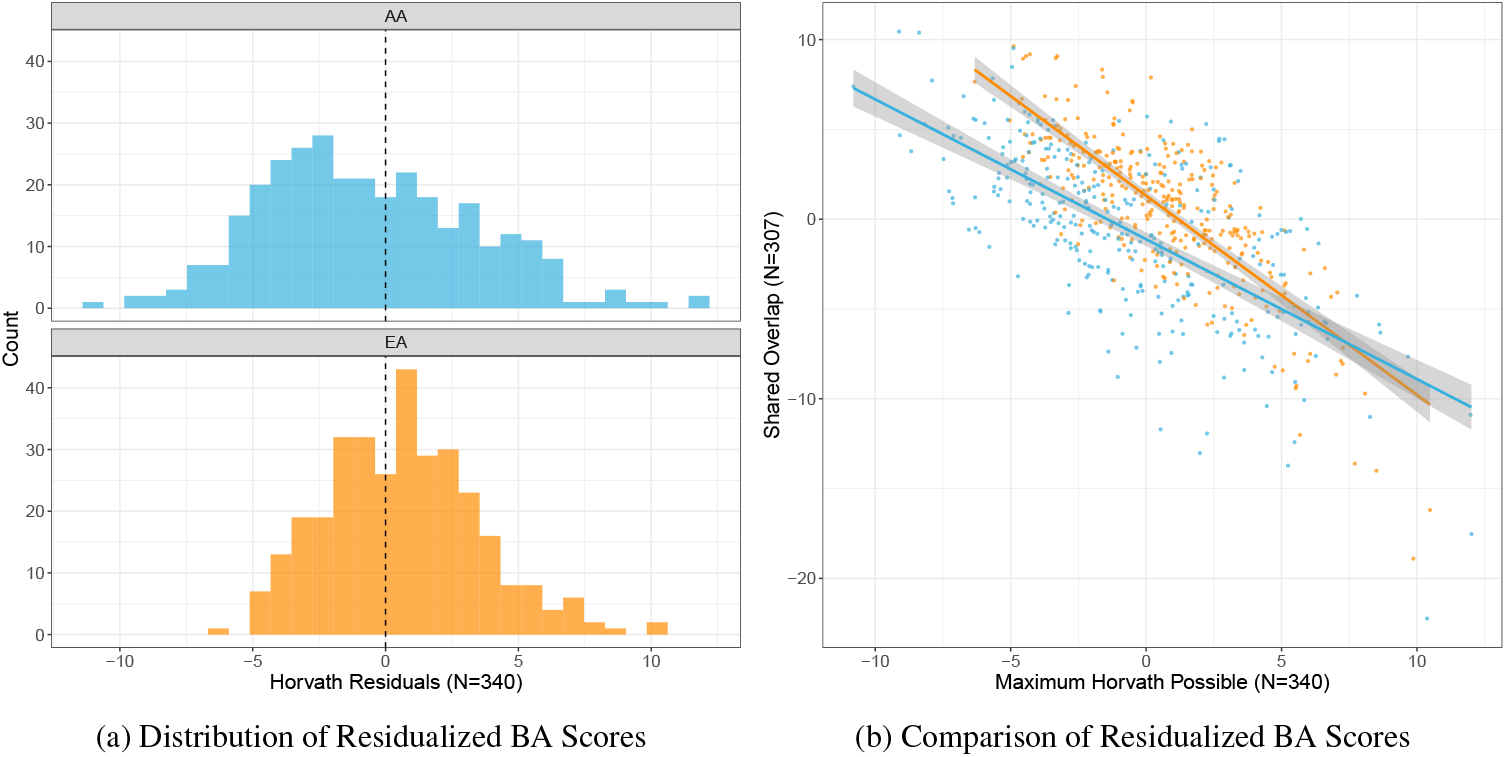
Residualized biological age scores were calculated by regressing chronological age on biological age (BA). (**3a**) The distribution of residualized BA scores by self-reported Census-based race category (AA = African American; EA = European American). (**3b**) Comparison of residualized BA scores calculated from the largest possible Horvath probe subset and the probe subset shared between PREG and replication cohort GAPPS (orange= EA, blue= AA).

## 4 Discussion

Excitement over the potential benefits associated with using BA to index personal risk liability for adverse health outcomes has prompted dozens of studies. ^7^ Indeed, such a biological marker could improve the accuracy of screening algorithms for multifactorial disorders. To our knowledge, this study is the first to examine the relationship between longitudinal measurements of prenatal maternal BA and GAAD. The results of this study highlight both potential benefits and caveats associated with using BA in translational research and clinical applications. Several characteristics of maternal prenatal BA are appealing for future follow up studies assessing clinical utility. Importantly, early prenatal BA was the most strongly associated with GAAD, which means that PTB risk assessments could occur in time to consider medical interventions and preventative measures. Further, this study observed large interindividual variation in baseline BA estimates which remained relatively stable throughout pregnancy. Early prenatal BA was significantly associated with GAAD above and beyond other risk factors like maternal prenatal perceived stress and chronological age. These findings suggest that early prenatal BA may be a promising candidate for inclusion in a precision clinical obstetrics screening algorithm.

Although results from this study support the possibility of adopting BA for estimating risk for PTB, some critical observations also were noted. First, sensitivity analyses revealed that the relationship between early prenatal BA and GAAD was impacted by probe set composition. Based on these findings, researchers should take care when estimating BA and clearly report the number of probes used in BA calculations. Second, the strongest association signal was found in the AA subset of the PREG sample. Although this relationship remained significant in the full PREG cohort after adjusting for self-identified Census-based race and multiple testing correction, sensitivity analyses using residualized BA scores suggest that the reliability of BA may vary by genetic ancestry and/or demographic factors. These findings suggest that cryptic, currently unidentified factors may be influencing the predictive validity and reliability of DNAm-based BA estimation. The problem of genomically-informed risk assessments failing to generalize to non-European populations has received increasing attention not only because such results limit the utility of clinical assessments but also because they threaten to exacerbate existing racial health disparities. ^42^ Another issue is that the biological significance of the individual sites of DNAm included in BA algorithms is poorly understood, ^43,44^ which obscures identifying the specific molecular processes BA actually reflects. ^7^ This knowledge gap makes predicting factors that will influence generalizability challenging. Researchers must be careful when studying populations that include individuals from a wide range of genetic ancestral backgrounds, especially given that most DNAm-based BA estimation algorithms work analogously to other methods that exhibit variable predictive validity by genetic ancestry (i.e., polygenic risk score calculation). ^42^

Although significant relationships were identified, the direction of the relationship between BA and GAAD was unexpected. Advanced biological aging is a putative driver of increased risk for negative health outcomes and would be expected in individuals with higher levels of perceived stress and pregnancies with a lower GAAD. In this case, the algorithm predicts that, on average, AA participants are biologically younger than their EA counterparts despite group differences in lifetime exposure to stressors that would predict greater positive deviations from chronological age. Given that a younger BA is associated with adverse outcomes during pregnancy, the results from this study may not support the traditional weathering hypothesis. The interpretation of BA-disease relationships may be complicated by the fact that risk for PTB is increased among both the youngest and oldest mothers, ^45^ rather than increasing over the lifetime like other age-related disorders. This nonlinear distribution between maternal chronological age and PTB may be similarly reflected in BA, so that any prominent deviations from mean BA, rather than advanced BA alone, may highlight those pregnancies at higher risk.

These findings contradict results from another study, which did not find a significant relationship between Horvath BA and GAAD, but did identify an inverse relationship between maternal BA estimated using another DNAm-derived BA algorithm and length of gestation. ^46^ However, other studies have similarly noted an unexpected direction of the association between DNAm-based BA and adverse pregnancy outcomes, including research assessing the relationships between the BA of infants at birth and maternal antenatal depression, PTB, and future psychiatric problems. ^47^ Contradictory relationships between fetal and placental telomere length, an alternative measure of cellular aging, and GAAD are also prevalent in the literature. ^32,31,48^ These results could arise from measurement variance that leads to unreliable BA estimates due to genetic and/or physiological status (i.e., pregnancy). The generalizability and reliability of genomic risk scores depends on the diversity and size of the training dataset composition, respectively. To our knowledge, no existing BA algorithm includes blood samples from pregnant women. As a result, BA estimates could be influenced by pregnancy-related DNAm remodeling. As the epigenetic aging field advances, BA estimators for specific populations have been developed, ^28,49,50^ and the development of future algorithms should be tailored for birth outcomes research and include pregnant women. Integrating DNAm-derived BA with other indices of cellular senescence (e.g., telomere length) could further increase our understanding of the molecular processes reflected in BA.

Overall, these results suggest that BA estimates hold potential to serve as a biomarker for PTB, but extreme care must be taken to assess the accuracy and generalizability of BA across a wide variety of genetic and demographic backgrounds. The ability to assess risk for PTB at the beginning of pregnancy would provide opportunities for early intervention and targeted medical care throughout gestation. Logistically, many attributes of DNAm-based BA make for a good candidate biomarker. ^51,52^ DNAm is a stable mark that can be measured reliably, and BA estimates are easily calculated using the Horvath method. In this study, DNAm was measured in peripheral blood, a tissue with a minimally invasive collection procedure that is already a normal part of pregnancy monitoring, posing no additional risk to patients. While more research is necessary to examine how reliably BA predicts GAAD in other samples, in the future BA should be considered for potential clinical applications.

### Strengths and limitations

To our knowledge, this study is the largest study to investigate maternal BA during pregnancy and is the first to examine the stability of prenatal BA and its relationship across time with GAAD. Major strengths of this study include the use of both a primary and replication cohort both containing longitudinal measurements during pregnancy. The inclusion of a diverse cohort allowed for the investigation of BA differences by self-reported race. Finally, all analyses and hypotheses examined in this study were preregistered on the Open Science Framework ^53^ using the AsPredicted format.

The results of this study should be considered in the context of four primary study limitations. One, cross-study comparisons were complicated by variation in data collection protocols. Perceived stress was assessed at four study visits in PREG while only two measures were collected in GAPPS. This limitation would have been easier to resolve if more detailed information about GA at assessment were available for GAPPS participants (e.g., GA in days). Two, the two study populations differed significantly in demographic composition (Table 1). These differences were particularly problematic given that main effects of BA on GAAD were seen primarily in the PREG AA subsample. Three, neither the PREG nor the GAPPS samples had complete probe data for the full Horvath algorithm. The GAPPS sample was measured using a newer technology missing seventeen of the Horvath probes, and both samples had probes removed during quality control. It is not clear if and how these missing probes influenced the final results, but the strength of the association between early prenatal BA and GA was attenuated in the smaller probe subset (n = 307) compared to the maximum possible probe subset for PREG (n = 340; see supplement for results from analyses including all available probes).

## 5 Additional information

## 5.1 Acknowledgements

The Pregnancy, Race, Environment, Genes (PREG) longitudinal study and its postpartum extension were supported by the NIMHD (P60MD002256, PI: York, Strauss), The John and Polly Sparks Foundation and Brain and Behavior Research Foundation (24712, PI: York) supported the use of GAPPS data and materials. Support was received from the Clinical and Translational Science Award (CTSA) award No. UL1TR000058 from the National Center for Advancing Translational Sciences. Its contents are solely the responsibility of the authors and do not necessarily represent official views of the National Center for Advancing Translational Sciences or the National Institutes of Health. Timothy York, Ph.D., holds a 2015 Preterm Birth Research Grant from the Burroughs Wellcome Fund. DML was supported by a T32 NIMH (PI: Michael Neale).

## 5.2 Preregistration

Analyses presented in this manuscript were preregistered on the Open Science Framework and are available at https://osf.io/6a9db. All of the original preregistered study questions were addressed in these analyses. However, there are other notable deviations from the analyses outlined in the preregistration document. Originally, two BA algorithms prominently featured in the literature, the Horvath and Hannum methods, were selected for this study. However, several probes included in the Hannum algorithm were removed during quality control processing steps. The Hannum method is known to be more sensitive to missing probes, potentially leading to a biased BA estimates. ^35^ As per the original preregistered study design, the same analyses were completed with the Hannum clock, (see supplement for relevant methods and results). Interestingly, both epigenetic clocks performed similarly in these samples, suggesting they are capturing the same biological phenomenon. Additionally, methods and results for a secondary analyses examining the use of chromosome Y probes to detect cell-free DNA contamination of maternal samples are available in the supplement. Finally, a more parsimonious model was selected to adjust for chronological age variability in the models. Rather than adopting a two-step approach in which BA is first regressed on chronological age before modeling the resulting residual, maternal age was simply included as a covariate in all analyses.

## 5.3 Data sharing

The preregistration document and R code used to analyze the data and generate figures is available on the Open Science Framework (OSF) project landing page (https://osf.io/sqmzg). Sharing PREG and GAPPS study data is limited by Institutional Review Board agreements and participant consent forms, which restrict openly sharing individual-level DNAm measures. Anyone interested in data access or collaboration is encouraged to contact Dr. Timothy P. York for more information.

## 5.4 Informed consent and ethical approvals

The PREG study received Virginia Commonwealth University Institutional Review Board approval (14000) and obtained written participant consent for each participant.

## 5.5 Author contributions

All authors designed the study. DL and EL cleaned the data, performed the analyses, and wrote the initial draft of the manuscript. CJC oversaw the processing/preparation of specimens for assessment in the PREG study. RRN, JS and TY provided substantive feedback and revisions, and TY and JS planned and secured funding for the PREG and GAPPS studies. All authors approved the final version of the manuscript.

## 5.7 Abbreviations

AA: African American
BA: Biological age
DNAm: DNA methylation EA European American
GAPPS: Global Alliance to Prevent Prematurity and Stillbirth
GA: Gestational age
GAAD: Gestational age at delivery
PREG: Pregnancy, Race, Environment, Genes study
PSS: Perceived Stress Scale
PTB: Preterm birth

