## Supplemental Materials for "Maternal Biological Age Assessed in Early Pregnancy is Associated with Gestational Age at Birth"

#### 1 Additional Demographic Information

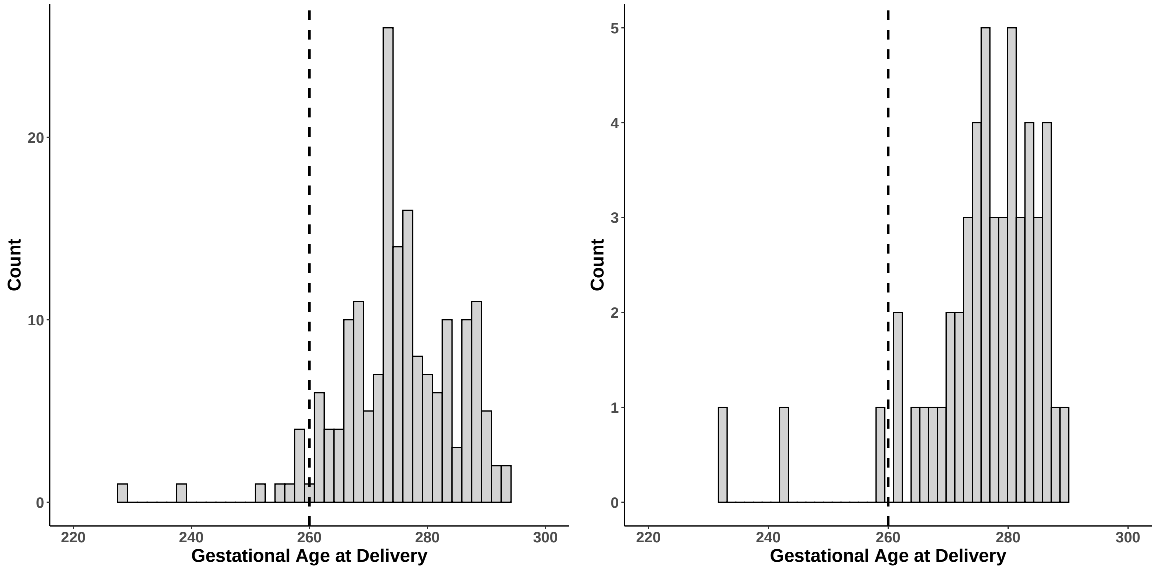


(a) Distribution of PREG GAAD

(b) Distribution of GAPPS GAAD

Figure S1: Distribution of gestational age at delivery across cohorts. (**S1a**) The distribution of gestational age at delivery (GAAD) in the full PREG cohort. (**S1b**) The distribution of GAAD in the GAPPS cohort.

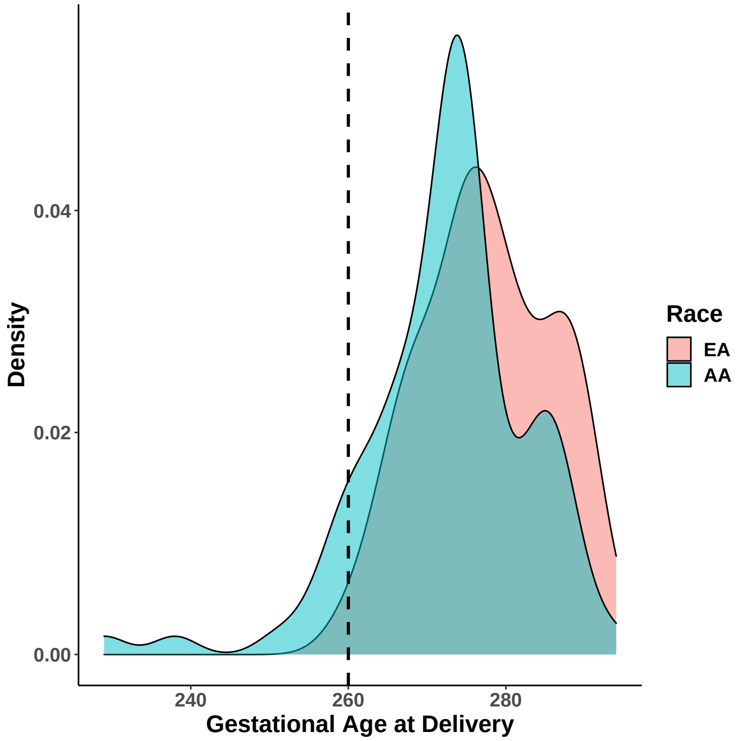


Figure S2: Distribution of PREG gestational age at delivery by race. The distribution of gestational age at delivery (GAAD) by self-reported Census-based race category (AA = African American; EA = European American).

### 2 Molecular measurements and raw data processing

All data processing steps were performed separately for each cohort using Bioconductor packages in the R programming environment.[1]

#### 2.1 DNA methylation measurement

Steps to generate raw measures of DNA methylation (DNAm) by microarray typically involve collecting tissue samples, extracting the DNA from those cells, performing sodium bisulfite conversion, hybridizing to the array, and reading the proportion of methylated to unmethylated signals. To complete these steps, whole blood was first collected from participants into EDTA tubes. Genomic DNA was isolated from 10 mL whole blood according to standard methods using the Puregene DNA Isolation Kit (Qiagen; Valencia, CA). An aliquot of 1 $\mu$g DNA per participant was sent for bisulfite conversion (HudsonAlpha Institute for Biotechnology; Zymo Research EZ Methylation Kit). Sodium bisulfite application deaminates an unmethylated cytosine residue to uracil, while 5-methylcytosine remains unaffected. This conversion step is critical for determining whether a given cytosine is methylated or unmethylated.

Genome-wide methylation was assayed according to the manufacturer's protocol at the HudsonAlpha Institute for Biotechnology (Illumina, San Diego, CA, USA). All specimens belonging to a single individual were placed on the same array to minimize the impact of array effects. Otherwise, samples were randomized to minimize any potential batch effects that might arise from array structure (i.e., differences by array, row, or column).

#### 2.2 DNA methylation preprocessing

Raw Intensity Data files (*.idat) files were generated after scanning the microarray chips and read into R for processing.[2] During standard DNAm preprocessing procedures, poor quality probes and samples are identified, data is normalized, and other potentially problematic probes are filtered. High detection p-values often indicate technological issues with the scanning of arrays (e.g., spatial artifacts). Probes with detection p-values greater than 0.01 were considered poor, and any probes that failed in more than 1% of samples were removed. Poor quality samples were defined as having both methylated and unmethylated median signal intensities less than 10.5 units and removed. Visual inspection confirmed that outlier samples had been eliminated.

Quantile normalization was performed to adjust for background correction, color bias, and probe type bias.[2,3] In addition to identifying those poorly performing probes, several other potentially confounding probes were removed, including cross-hybridizing probes and those probes with polymorphisms at the CpG site and single-base extension. Finally, Beta-values were further adjusted for slide-related technical artifacts prior to analysis using an empirical Bayes framework.[4]

### 3 Adjusting for missing probes

In the initial preregistration document, we did not consider the possibility that relevant DNAm probes could be missing from the PREG study. We expected some probes to be non-overlapping between the PREG and GAPPS samples due to differences in beadchip technology; however, we were not concerned by this issue because the 17 missing probes had little impact on biological age estimate correlations in samples assayed with both the Illumina EPIC and HumanMethylation450 microarrays.[5,6]

**PREG Study.** In the PREG study, we identified 13 probes missing from the Horvath probe set (3.7%). The combined effect size of the missing probe set was +0.27. Given that the missing probes were relatively balanced for positive and negative influences, we felt comfortable proceeding with the Horvath age estimates and residuals calculated for the PREG study.[7]

**GAPPS Study.** In the GAPPS study, we identified 33 probes missing from the Horvath probe set (9.3%). The combined effect size of the missing probe set was -0.48 years. Given that the missing probes were relatively balanced for positive and negative effect sizes, we felt comfortable proceeding with the Horvath age estimates and residuals calculated for the GAPPS study.

**Shared Probe Analysis.** In the PREG study, we identified 307 probes present in both the PREG and GAPPS samples for the Horvath algorithm (87%). Given that the missing probes were relatively balanced for positive and negative effect sizes, we felt comfortable proceeding with the Horvath age estimates and residuals calculated for the GAPPS study.

Overall results were consistent whether biological age was estimated by the maximum amount of Horvath probes per cohort or using only probes shared between PREG and GAPPS. A statistically significant relationship between early-pregnancy biological age and gestational age at delivery remained after adjusting for the six within-cohort tests using the Bonferroni method ($p < 0.008$) in the full PREG sample only (Table S1 and S2). A significant relationship was also identified between early biological age and perceived stress during early pregnancy in the full PREG sample and in the African American subset.

Table S1: Relationships of Gestational Age at Delivery, Perceived Stress, and Horvath Biological Age Estimates Calculated Using All Available Probes in PREG[1]

| | PREG Coef | PREG SE | PREG P-value[2] | PREG AA Coef | PREG AA SE | PREG AA P-value[2] | PREG EA Coef | PREG EA SE | PREG EA P-value[2] |
|---|---|---|---|---|---|---|---|---|---|
| Predicts GAAD | | | | | | | | | |
| early BA | 0.607 | 0.213 | 0.005* | 0.682 | 0.352 | 0.059 | 0.374 | 0.239 | 0.124 |
| late BA | 0.115 | 0.172 | 0.503 | 0.090 | 0.241 | 0.708 | -0.029 | 0.254 | 0.910 |
| Predicts early prenatal PSS | | | | | | | | | |
| early BA | -0.390 | 0.131 | 0.004* | -0.506 | 0.179 | 0.007* | -0.179 | 0.182 | 0.330 |
| Predicts late prenatal PSS | | | | | | | | | |
| early BA | -0.165 | 0.147 | 0.265 | -0.162 | 0.210 | 0.447 | -0.126 | 0.203 | 0.538 |
| late BA | -0.091 | 0.121 | 0.453 | -0.013 | 0.150 | 0.933 | -0.021 | 0.195 | 0.915 |
| Predicts late BA | | | | | | | | | |
| early PSS | -0.085 | 0.051 | 0.099 | -0.049 | 0.087 | 0.571 | -0.050 | 0.066 | 0.451 |

[1] The maximum number of probes were retained to most accurately match the original Horvath probe set.

[*] Survives Bonferroni adjustment $p < 0.008$

**Abbreviations.** Coef = coefficient; SE = standard error; EA = European-American; AA = African-American; PSS = perceived stress scale total score; GAAD = gestational age at delivery.

Table S2: Relationships of Gestational Age at Delivery, Perceived Stress, and Horvath Biological Age Estimates Calculated Using All Available Probes in GAPPS[1]

| | GAPPS Coef | GAPPS SE | GAPPS P-value[2] |
|---|---|---|---|
| Predicts GAAD | | | |
| early BA | -0.114 | 0.501 | 0.821 |
| late BA | 0.372 | 0.412 | 0.371 |
| Predicts early prenatal PSS | | | |
| early BA | -0.226 | 0.234 | 0.341 |
| Predicts late prenatal PSS | | | |
| early BA | -0.394 | 0.258 | 0.134 |
| late BA | -0.140 | 0.258 | 0.592 |
| Predicts late BA | | | |
| early PSS | -0.014 | 0.104 | 0.898 |

[1] The maximum number of probes were retained to most accurately match the original Horvath probe set.

[*] Survives Bonferroni adjustment $p < 0.008$

**Abbreviations.** Coef = coefficient; SE = standard error; PSS = perceived stress scale total score; GAAD = gestational age at delivery.

### 4 Growth Models

The purpose of the growth curve model was to quantify the stability and rate of change of residualized biological age estimates across pregnancy and to test if a relationship existed between the growth parameters and gestational age at delivery. Growth models were run using the `MplusAutomation` library. The longitudinal data came from the 177 PREG participants who met all inclusion and no exclusion criteria. The code for the models can be found on this project's Open Science Framework page (https://osf.io/sqmzg/). Briefly, models were built and run in order of complexity, starting with a simple linear growth model without covariates or a distal outcome. Time scores (i.e., TSCOREs) were used to adjust for interindividual variation in the timing of DNAm collection. Gestational age at assessment was divided by 100 to match the scale (i.e., mean and variance) of the residualized biological age estimates. Any model that converged but did not generate standard error estimates was considered untrustworthy and uninterpretable.

The unadjusted linear growth curve models (LGCM) with and without gestational age at delivery as a distal outcome converged. The unadjusted LGCM with a distal outcome suggested that both growth parameters predicted gestational age at delivery (intercept $p = 0.003$; and slope $p < 0.001$). Unfortunately, the interpretability of these models is uncertain because models with key covariates (e.g., maternal smoking status and self-identified Census-based race categories) failed to converge. As a result, the partial correlations for residualized biological age estimate, race, and smoking status could not be estimated. To overcome this problem, a new LGCM using the biological age estimate itself and a covariate for age was tested. Biological age estimates were divided by 10 and gestational age at assessment was divided by 100 to put the variables on comparable scales. This new LGCM converged when current maternal smoking status was added as a covariate, and the results suggested that smoking status was significantly associated with both the intercept and slope even without a distal outcome.

The trajectories of residualized biological age scores appeared to have significant between-person heterogeneity in terms of both initial measurements and the change in magnitude over the course of pregnancy. Within person, biological age either remained largely the same or decreased a small amount towards the end of pregnancy. Given the significant technical issues associated with the LGCMs, no firm conclusions can be drawn about how the trajectories of maternal biological age estimates or residualized biological age estimates associate with gestational age at delivery. Future studies should not assume high maternal biological age stability across pregnancy. More work is needed to know if the stability observed in this study is generalizable. All of the PREG participants used for this analysis had either full term or late preterm deliveries of healthy babies. It is possible that maternal biological age during pregnancy behaves very differently when pregnancy complications like preeclampsia or RH sensitization are present.

### 5 Deviations from Preregistration

#### 5.1 Associations between Hannum-derived biological age estimates and pregnancy outcomes

Originally, two biological age algorithms prominently featured in the literature, the Horvath and Hannum methods, were selected for this study. However, several probes included in the Hannum algorithm were removed during quality control processing steps. The large combined effect size of the removed probes, alongside reports that the Hannum method is known to be more sensitive to missing probes,[6] lead to concerns about focusing on the potentially biased Hannum biological age estimates.

**PREG Study.** In the PREG study, we identified four probes missing from the Hannum probe set (5.6%). The combined effect size of the missing probes was +20 years. Given the large bias in the estimates of the missing probes, we decided not to proceed with using Hannum-derived biological age estimates as a primary analysis in the PREG cohort.[8]

**GAPPS Study.** In the GAPPS study, we identified 10 probes missing from the Hannum probe set (14.1%). The combined effect of the missing probes was +23.28 years. Given the large bias in the estimates of the missing probes, we decided not to focus on Hannum-derived biological age estimates in the GAPPS sample.

Given that neither cohort had the full probeset available, biological age estimates were calculated both with all available probes and with only those probes remaining in both cohorts. Both of the resulting estimates were separately tested for relationships with gestational age at delivery and perceived stress during pregnancy.

A statistically significant relationship between early-pregnancy biological age and gestational age at delivery remained after adjusting for the six within-cohort tests using the Bonferroni method ($p < 0.008$) in the full PREG sample only (Table S3 and S4). A significant relationship was also identified between early biological age and perceived stress during early pregnancy in the full PREG sample. When calculating biological age using only those Hannum probes that were present in both samples, no relationships reached significance (Tables S5 and S6). Overall, biological age estimated using the Hannum algorithm showed similar relationships with the tested outcomes as Horvath-estimated biological age. Early biological age was more predictive, and the direction of the relationships were again unexpected (younger biological age associated with higher perceived stress and shorter gestational age at delivery) based on a weathering hypothesis.

Table S3: Relationships of Gestational Age at Delivery, Perceived Stress, and Hannum Biological Age Estimates Calculated Using All Available Probes in PREG[1]

| | PREG Coef | PREG SE | PREG P-value[2] | PREG AA Coef | PREG AA SE | PREG AA P-value[2] | PREG EA Coef | PREG EA SE | PREG EA P-value[2] |
|---|---|---|---|---|---|---|---|---|---|
| Predicts GAAD | | | | | | | | | |
| early Hannum | 0.771 | 0.274 | 0.006* | 0.739 | 0.398 | 0.070 | 0.509 | 0.368 | 0.173 |
| late Hannum | 0.380 | 0.184 | 0.041 | 0.432 | 0.307 | 0.164 | 0.236 | 0.228 | 0.305 |
| Predicts early prenatal PSS | | | | | | | | | |
| early Hannum | -0.617 | 0.163 | 0.000* | -0.662 | 0.195 | 0.002* | -0.340 | 0.278 | 0.228 |
| Predicts late prenatal PSS | | | | | | | | | |
| early Hannum | -0.483 | 0.202 | 0.019 | -0.539 | 0.247 | 0.037 | -0.348 | 0.321 | 0.284 |
| late Hannum | -0.042 | 0.157 | 0.789 | 0.129 | 0.216 | 0.551 | -0.029 | 0.222 | 0.897 |
| Predicts late Hannum | | | | | | | | | |
| early PSS | -0.070 | 0.047 | 0.140 | -0.052 | 0.066 | 0.433 | -0.039 | 0.073 | 0.591 |

[1] The maximum number of probes were retained to most accurately match the original Hannum probe set.

* Survives Bonferroni adjustment $p < 0.008$

**Abbreviations.** Coef = coefficient; SE = standard error; EA = European-American; AA = African-American; PSS = perceived stress scale total score; GAAD = gestational age at delivery.

Table S4: Relationships of Gestational Age at Delivery, Perceived Stress, and Hannum Biological Age Estimates Calculated Using All Available Probes in GAPPS[1]

| | GAPPS Coef | GAPPS SE | GAPPS P-value[2] |
|---|---|---|---|
| Predicts GAAD | | | |
| early Hannum | -0.931 | 0.758 | 0.226 |
| late Hannum | 0.511 | 0.526 | 0.336 |
| Predicts early prenatal PSS | | | |
| early Hannum | -0.377 | 0.365 | 0.308 |
| Predicts late prenatal PSS | | | |
| early Hannum | -0.704 | 0.393 | 0.080 |
| late Hannum | -0.581 | 0.320 | 0.076 |
| Predicts late Hannum | | | |
| early PSS | -0.096 | 0.082 | 0.249 |

[1] The maximum number of probes were retained to most accurately match the original Hannum probe set.

[*] Survives Bonferroni adjustment $p < 0.008$

**Abbreviations.** Coef = coefficient; SE = standard error; PSS = perceived stress scale total score; GAAD = gestational age at delivery.

Table S5: Relationships of Gestational Age at Delivery, Perceived Stress, and Hannum Biological Age Estimates Calculated Using Overlapping Probes in PREG[1]

| | PREG Coef | PREG SE | PREG P-value[2] | PREG AA Coef | PREG AA SE | PREG AA P-value[2] | PREG EA Coef | PREG EA SE | PREG EA P-value[2] |
|---|---|---|---|---|---|---|---|---|---|
| Predicts GAAD | | | | | | | | | |
| early Hannum | 2.300 | 1.650 | 0.167 | 2.348 | 2.584 | 0.369 | 1.358 | 1.904 | 0.479 |
| late Hannum | -1.245 | 1.041 | 0.233 | -1.288 | 1.473 | 0.385 | -1.806 | 1.462 | 0.220 |
| Predicts early prenatal PSS | | | | | | | | | |
| early Hannum | -1.779 | 1.009 | 0.081 | -2.792 | 1.316 | 0.040 | -0.062 | 1.440 | 0.966 |
| Predicts late prenatal PSS | | | | | | | | | |
| early Hannum | -0.562 | 1.149 | 0.626 | -0.596 | 1.482 | 0.691 | -0.449 | 1.702 | 0.793 |
| late Hannum | 0.471 | 0.765 | 0.539 | 0.558 | 0.948 | 0.559 | 1.113 | 1.188 | 0.352 |
| Predicts late Hannum | | | | | | | | | |
| early PSS | 0.003 | 0.008 | 0.755 | 0.001 | 0.014 | 0.939 | 0.012 | 0.011 | 0.311 |

[1] Hannum probe sets were reduced to match the probes available for GAPPS.

[*] Survives Bonferroni adjustment $p < 0.008$

**Abbreviations.** Coef = coefficient; SE = standard error; EA = European-American; AA = African-American; PSS = perceived stress scale total score; GAAD = gestational age at delivery.

Table S6: Relationships of Gestational Age at Delivery, Perceived Stress, and Hannum Biological Age Estimates Calculated Using Overlapping Probes in GAPPS[1]

| | GAPPS Coef | GAPPS SE | GAPPS P-value[2] |
|---|---|---|---|
| Predicts GAAD | | | |
| early Hannum | -0.914 | 0.792 | 0.255 |
| late Hannum | 0.531 | 0.532 | 0.323 |
| Predicts early prenatal PSS | | | |
| early Hannum | -0.425 | 0.380 | 0.270 |
| Predicts late prenatal PSS | | | |
| early Hannum | -0.784 | 0.407 | 0.061 |
| late Hannum | -0.599 | 0.323 | 0.070 |
| Predicts late Hannum | | | |
| early PSS | -0.100 | 0.081 | 0.224 |

[1] Hannum probe sets were reduced to match the probes available for GAPPS.

[*] Survives Bonferroni adjustment $p < 0.008$

**Abbreviations.** Coef = coefficient; SE = standard error; PSS = perceived stress scale total score; GAAD = gestational age at delivery.

#### 5.2 Chromosome Y Analyses

The preregistration describes a sensitivity analysis intended to investigate the potential influence of cell-free fetal DNA (cffDNA) and DNAm on prenatal maternal DNAm-based biological age estimates. Given that pregnant mothers typically do not carry a copy of chromosome Y (chrY), probes on chrY were used as a proxy measure of cffDNA.

**Motivation.** Pregnant women have measureable levels of cffDNA in their peripheral blood that can be used to detect aneuploidy and sex chromosome abnormalities reliably. It is unknown the extent that cffDNA influences DNAm measurement in maternal blood. The 450k beadchip uses 50bp probes to interrogate the DNA methylome. During sample preprocessing, DNA is sheared into 150bp-200bp fragments. The average length of cffDNA is 160 bp and has been measured $>$340bp, so the fragments on average have enough length to be probed.[9] One key observation by Fan et al. (2010) was that the average length of chrY fragments was shorter than other chromosomes ("rarely longer than 250bp and present at $<$150bp").

**Methods.** Female specimens from the Adolescent and Young Adult Twin Study (AYATS) were used to identify chrY probes with cross-reactivity (i.e., had measurable signal in a never-pregnant sample of adolescent females). The signal intensity of chrY probes was compared to signal profiles from other chromosomes and from control probes in a mixed-sex subset of AYATS participants ($n = 133$). Reliable chrY probes were then used in PREG participants who are carrying male children to evaluate the level of chrY probe signal. Both single probe and multiple probe approaches were tested for a subset of chrY probes.

**Results.** The copy number analysis showed that both chromosome X and chrY copy numbers were low compared to small autosomes like chromosome 20. The level of chrY signals appeared to be similar to background ground levels of the control probes. The chrY signal almost entirely vanishes when males are removed. Most chrY probes do not produce signal intensities that exceed the highest quantile of negative control probe signal intensity in an all female sample of never-pregnant adolescents. The probe(s) with the most signals over the highest negative control signal were cg04964672 and cg01086462, which are both located in the TTTY16 gene on short arm of the chrY. It is possible that this elevated signal is due to cross-hybridization, aberrant tiling (i.e., the probes are not lying down on the DNA fragments correctly or have been mis-annotated), or polymorphisms in the pseudoautosomal region.

Neither single nor multiple probe approaches predicted gestational age at birth or current gestational age in the PREG sample. The pattern of chrY signal was erratic. Even proximal probes showed low correlations ($r \sim .10$), and this pattern held for the SRY region on chrY. The correlations of chrY probes were nominally higher in maternal blood samples from women pregnant with male offspring; however, if an effect is present, it is very small.
